# Synthesising integrated robot behaviour through reinforcement learning for homeostasis

**DOI:** 10.1101/2024.06.03.597087

**Authors:** Naoto Yoshida, Hoshinori Kanazawa, Yasuo Kuniyoshi

## Abstract

Homeostasis is a fundamental property for the survival of animals. Computational reinforcement learning provides a theoretically sound framework for learning autonomous agents. However, the definition of a unified motivational signal (i.e., reward) for integrated survival behaviours has been largely underexplored. Here, we present a novel neuroscience-inspired algorithm for synthesising robot survival behaviour without the need for complicated reward design and external feedback. Our agent, the *Embodied Neural Homeostat*, was trained solely with feedback generated by its internal physical state and optimised its behaviour to stabilise these internal states: homeostasis. To demonstrate the effectiveness of our concept, we trained the agent in a simulated mechano-thermal environment and tested it in a real robot. We observed the synthesis of integrated behaviours, including walking, navigating to food, resting to cool down the motors, and shivering to warm up the motors, through the joint optimisation for thermal and energy homeostasis. The Embodied Neural Homeostat successfully achieved homeostasis-based integrated behaviour synthesis, which has not previously been accomplished at the motor control level. This demonstrates that homeostasis can be a motivating principle for integrated behaviour generation in robots and can also elucidate the behavioural principles of living organisms.

---

For survival, mammals and other animals strictly regulate the internal conditions of their bodies within specific ranges. Homeostasis, a fundamental concept in physiology and the autonomic nervous system ^1,2^, is also well recognized at the behavioural level ^3^. For example, animals eat to maintain energy levels for activity and shiver to regulate body temperature ^4,5^. These behaviours are seamlessly integrated into the animal’s movements.

But how is this possible?

To produce integrated behaviors in artificial agents, homeostasis has been proposed numerous times as a goal for the autonomous and generic emergence of behavior in both neuroscience and robotics ^6-8^. Previous studies typically assumed the presence of behavior modules, which were often handcrafted within models. These frameworks selected modules based on evaluation indices linked to homeostasis but did not account for the organism-like generation of new behaviors ^6,7^. However, it remains underexplored whether the principle of homeostasis can lead to behavior emergence at the motor control level in realistically configured artificial robotic systems.

In this study, we construct an autonomous real robot system where homeostasis serves as the sole learning objective. Our results demonstrate the possibility of the emergence of integrated behaviors aimed at maintaining homeostasis at the motor control level. The bio-inspired robot developed in this research represents the world’s first real robot system to achieve the emergence of integrated behaviors based solely on the principle of homeostasis.

In recent computational neuroscience, homeostatic reinforcement learning theory (HRL) has been proposed. This theory uses classical homeostatic regulation as a paradigm for behavioural learning through computational reinforcement learning ^9,10^. HRL is based on the natural assumption that animals have bodies, and by treating homeostasis as the fundamental optimization objective for behavioural learning, various adaptive animal behaviours can be explained as processes of behavioural optimization (Fig. 1a). However, in previous neuroscientific modelling and behavioural optimization contexts, this theory has been applied only to small-scale problems and has not addressed the emergence of complex behaviours ^10^.

**Fig. 1.**
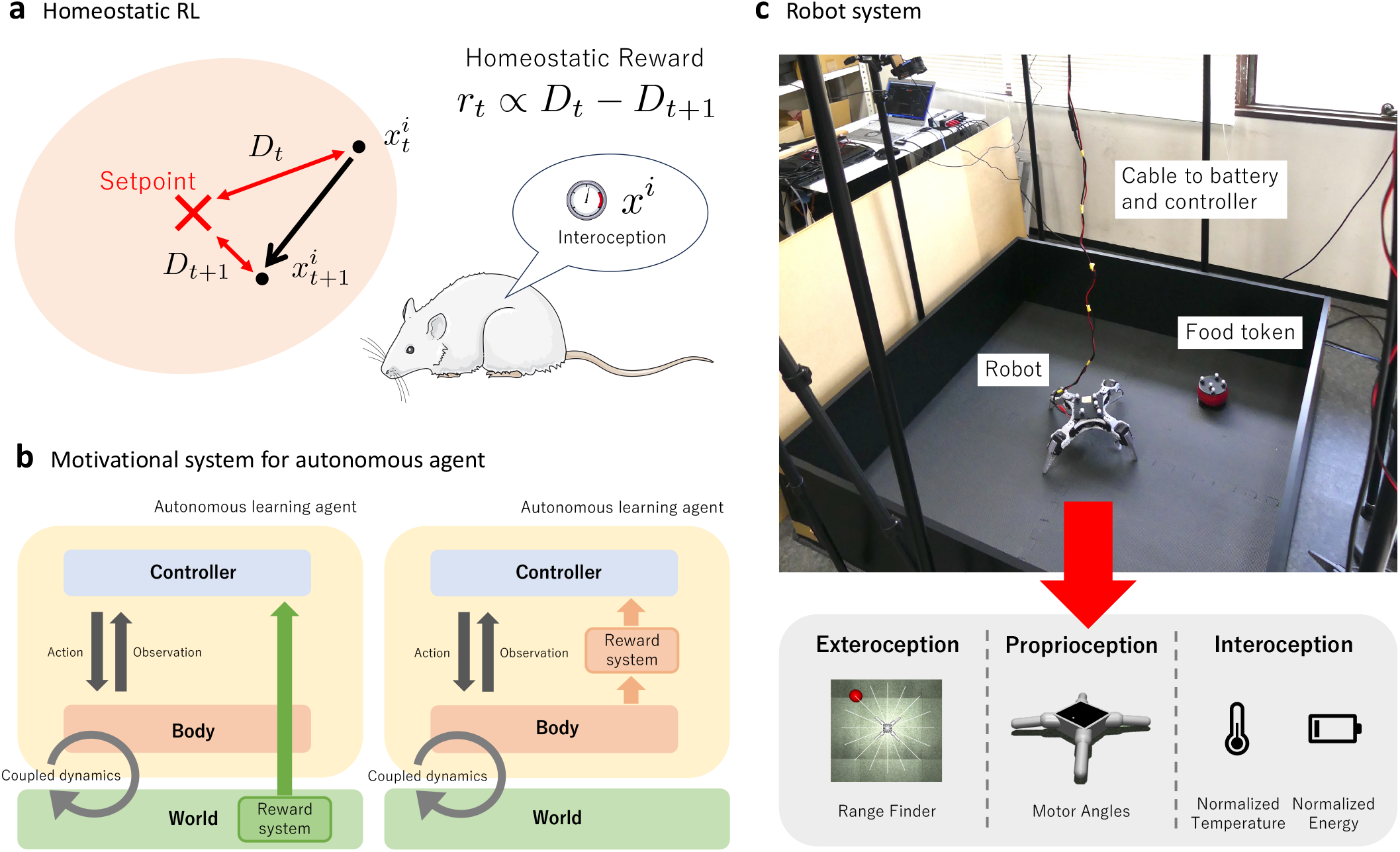
Configuration. **a**, Overview of homeostatic RL. In homeostatic RL, the agent has an internal state in its body and observes this state as an interoception *x*^*i*^. The interoception has a range of values (red colored region) and a target value (setpoint), and the accumulation of errors *D* between the observed interoception and the target value in the future is minimized. This is accomplished by defining the reward in terms of the difference in error between the two timesteps *r*_*t*_ ∝ *D*_*t*_ − *D*_*t* +1_. **b**, Reward systems as autonomous learning agents. Left: a typical RL setting where the agent is trained by an external reward system. Right: reward system as a motivational system internal to the body in a homeostatic RL agent. The coupled dynamics of the agent’s body and the external world allows the agent to manipulate the internal state of its own body directly and indirectly. For the agent to voluntarily acquire this control, a generic reward system closed inside the body is necessary in autonomous agents such as animals. **c**, Configuration of the robot system in this study (top), in which the robot body and a food token is placed in a square field, and the position of the food token is probabilistically placed by the experimenter at the position specified by the experiment management system. The bottom part of the figure shows multimodal observations of agents, based on Sherrington’s classification.

In recent years, deep reinforcement learning (deep RL) has demonstrated performance surpassing that of humans in various control tasks. Examples include Atari games ^11^, Go ^12^, StarCraft ^13^, Dota ^14^, Gran Turismo ^15^, and real-world drone racing ^16^. Deep RL has proven to be an effective machine learning method for these control tasks, where it is challenging for humans to generate large amounts of correct data.

In these previous studies, the reward configuration required during RL agent training is given by the agent’s external environment (Fig. 1b, Left). In other words, an external oracle-like evaluation system is assumed ^17^. This view of agent and environment ignores the rich interaction between the coupled internal physical dynamics of the body and the external environment ^18^, which is in fact present in natural animals and real robots (Fig. 1b, Right).

Homeostatic RL assumes the coupling of internal and external body dynamics and optimises behaviour so that the distance between the body’s internal state *x*^*i*^ and the target value is minimised over the future. (Fig. 1a). By extending Homeostatic RL with the acquisition of complex behaviours made possible by deep RL, attempts have been made to generate behaviours solely by rewards that originate from physiological dynamics computed from within the body ^19,20^. These studies have observed the emergence of behaviours such as foraging control of mobile robots with different nutritional states in simulation ^19,^ and, more recently, walking and temperature control through motor torque control of a simulated quadruped robot with simplified nutrient metabolism and thermal dynamics ^20^.

However, all these previous studies were attempts in simplified simulations, not in the real world by using robot with the power and temperature dynamics. Therefore, the possibility of behaviour emergence based on the principle of homeostatic RL on actual robotic systems, which can have complex internal interdependencies, remains to be verified.

In this article, we describe Embodied Neural Homeostat (ENH), a system for learning behaviours in real robots using only internal physical states of the body to define rewards. For the first time, this study offers evidence that integrated behaviours can emerge in real robot systems using only the homeostatic reward based on the internal physical states of the robot’s body.

The system situation in this study is shown in Fig. 1c: this is survival-demanding environments where robots need to feed themselves to control their own battery levels, and thermal regulation. A quadruped robot is placed inside a square field with walls. The robot can drive its motors using an externally connected battery. This battery can be replaced during the execution of the experiment and is recharged by being physically replaced when certain conditions are met, as described below. In addition to the robot, a food token (red object) is placed in the field. Both the robot and the food token are tracked by the tracking system, and the experiment is stopped when the robot detects a certain interaction with the food token, and the robot is recharged by performing a series of battery replacement processes. The robot has temperature sensors inside all its motors, and battery monitoring sensors constantly monitor the batteries to receive information. The purpose of the robot in this study is to control the robot’s body temperature measured by the temperature sensor and the battery level in this environment to establish a thermal homeostasis and an energy homeostasis at the same time.

## Approach

The controller of the ENH receives multimodal observation inputs from the environment, including the robot and battery, at approximately 20 Hz (Fig. 2). Here, referring to Sherrington’s classification^21^, observations of the outside of the robot’s body are referred to as exteroception, observations of its own posture, such as the position of joint angles, are referred to as proprioception, and observations of the internal state of the body are referred to as interoception.

**Fig. 2.**
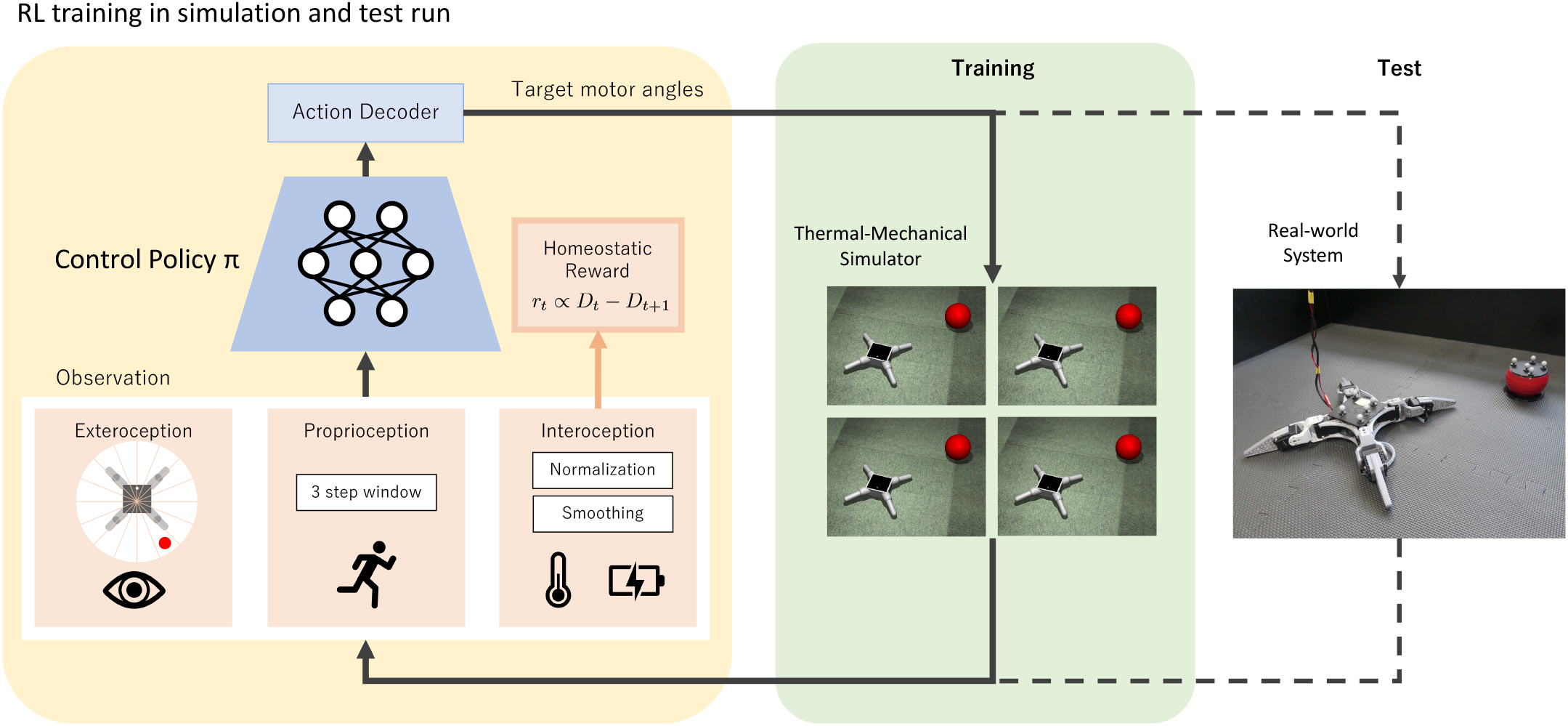
ENH training and test. Setup for RL. A deep neural network (control policy) receives multimodal information as input and outputs actions probabilistically. The output actions were converted into motor command values for the robot by the action decoder. All rewards during training were calculated from interoception, and rewards for homeostasis were generated. An environmental model created in advance by a thermal-mechanical simulator was used to train the network. The trained network was applied to a real robot system to configure a real-world ENH and evaluate its behaviour.

In the system configuration of the ENH in this study, the exteroception is a virtual rangefinder sensor that returns the distance to the prey, with a radial detection range centred on the robot. This was configured by obtaining the position and orientation of each of the food token and the robot from the tracking system data. This proprioception is an observation of the angular information of each of the robot’s eight axis joints stacked for three-time steps. Finally, the interoception consists of the robot’s body temperature and remaining energy. These are defined, respectively, by the body temperature as the average value after smoothing and normalizing the temperatures of all motors, and by the remaining energy as the normalized value of the amount of charge consumed since the battery was recharged.

The controller consists of a multilayer perceptron with two hidden layers, which outputs 9-dimensional continuous values. These consist of an 8-dimensional target value for the joint angles and a 1-dimensional probability. The latter probability represents the probability of outputting a joint angle that results in a specific cooling posture for the robot. These outputs are input to a pre-configured action decoder, which outputs the actual target angles to the 8-axis servo motors to the robot system (Extended Data 1).

The optimization procedure of ENH was built based on an on-policy model-free deep RL algorithm ^22^. We adopted a transfer learning approach from simulation to real robot and trained ENH only during simulation: the ENH controller is connected to the simulator and performs on-policy model-free deep RL optimization using sampled data of parallel interaction trajectories with multiple clones. In training, only rewards proportional to the time difference between the distance (drive) *D*_*t*_ between the robot’s interoception 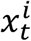 and its target value 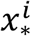 were used.

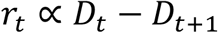

This is a reward setting that allows autonomous learning because it can be computed using only the information inside the robot system.

The simulation environment used for training consisted of a mechanical simulator and a lightweight thermal simulator, each of which was configured and run simultaneously. For this purpose, shape and motion data of the robot were collected, and the data were fitted to the simulators to make the motion and temperature changes in the real system virtually predictable. On the other hand, it has been pointed out that errors between simulation and reality can be problematic in real-world applications of optimised controllers ^23^. Since the robot geometry is modelled with a simplified structure in this study, the physical properties of the machine details and the nature of friction may differ from reality. Therefore, our optimization absorbed these hardware errors by randomly changing the mechanical properties of the simulation environment, including the robot. Furthermore, in the actual robot, there is a communication delay, so the simulation error may be caused by time delay as well. In a preliminary experiment, we compared the dynamics of the joint angles of the robot in the simulation with the actual joint angles. The delay in the real system was negligible within the range of our experimental system, and the time delay was ignored in this study.

After optimization, we verified the behaviour of ENH using a real robot system. To systematize this verification, the sequence of operation of the experiment was managed by a PC. This automated instructions on when to change batteries and where to place the food token, and the experimenter followed these instructions to recharge the robot and relocate the food token.

## Results

ENH can operate continuously in the environment by controlling body temperature and battery power through interaction with the environment.

Examples of long-term results from continuous operation of ENH in the field are shown in Fig. 3a, b; Fig. 3a shows the actual trajectory of internal states. This plot demonstrates that ENH can stably control the residual energy and body temperature around the target values (dashed line). It can be confirmed that homeostasis of the body’s internal state can be realized in the real robot over a long period of time.

**Fig. 3.**
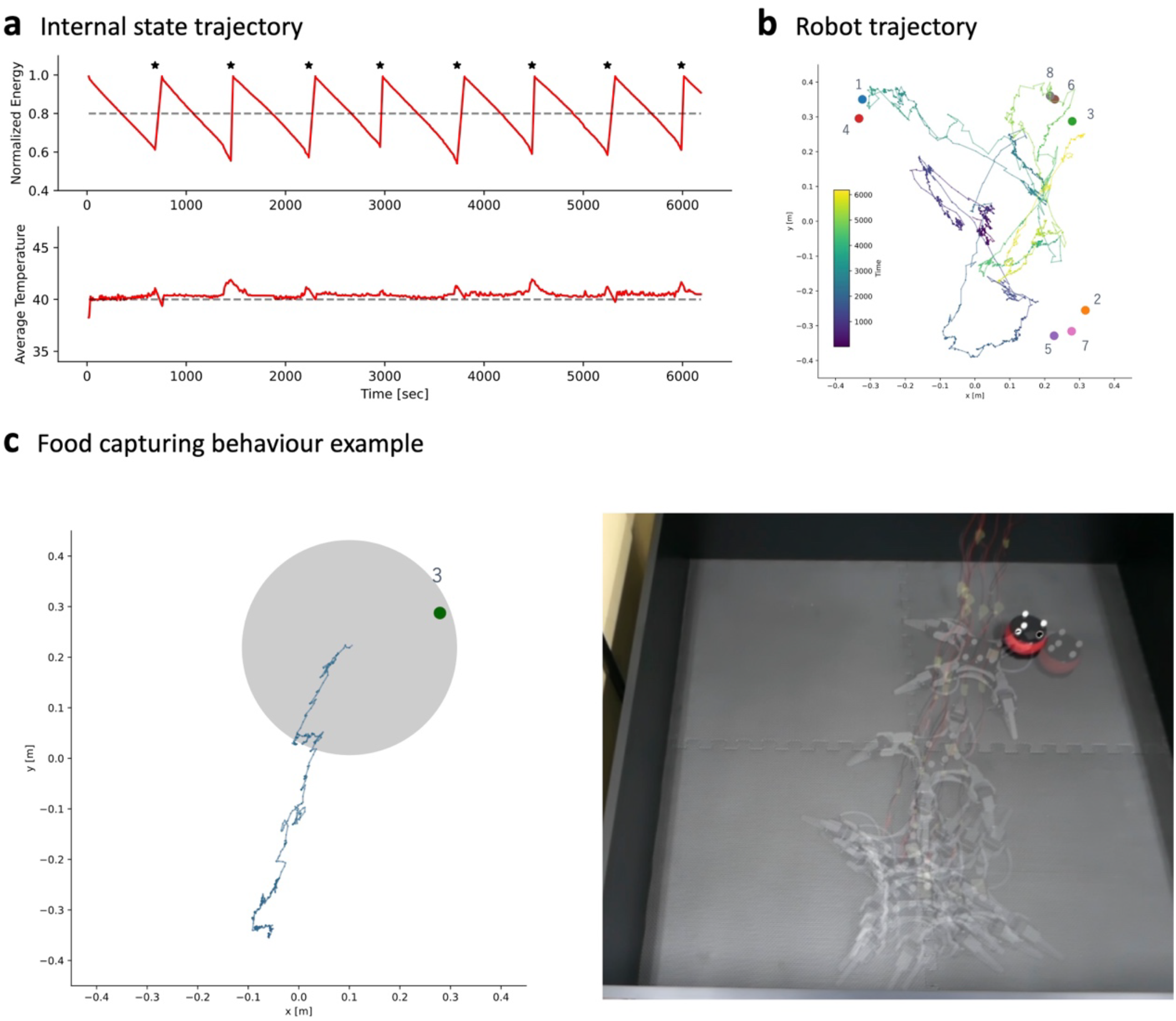
Results. **a,** Trajectories of internal states of trained ENHs in a real environment. The dashed lines represent the target value of each internal state. The star in the figure represents the timing at which the battery was reset by the robot’s interaction with the food token. **b**, The trajectory of ENH in the field during the experiment and the positions of the eight food tokens placed during the experiment. **c**, ENH during the 1,000 steps immediately before the intake of the third placed food token. trajectory (left) and the corresponding time-lapse image of the actual robot locomotion (right). The grey circles represent the approximate size of the robot’s legs spread out.

Observation of the behaviour of the ENH showed that it repeatedly waited at the same location in the field and moved to the food token when the battery level was low (Fig. 3b, Supplementary Video 1). Therefore, we can confirm that the ENH is very different from the static impression given by the term “homeostasis,” and that it moves considerably throughout the field to maintain its homeostasis. Fig. 3c shows a specific example of such movement. It shows the trajectory of ENH in the third situation in which it collected a food token (left), plus a time lapse of the robot at that time (right).

In principle, all ENH behaviours are integrated for homeostasis, and all movements are emergent, unlike approaches that train and integrate individual behaviours such as walking and navigation separately.

In an ablation study in which temperature homeostasis was not a learning target, the results showed that the ENH controlled energy while the body temperature continued to rise (Extended Data 3). This result suggests that body temperature control is essential for the long-term operation of ENH in the experimental setup of this study.

In addition, the actual battery replacement sequence temporarily shuts down the power supply of the robot. Therefore, the associated drop in body temperature may make it appear that ENH homeostasis has been achieved. As an ablation study to verify this possibility, we performed a similar experiment by creating a situation in which the energy level was reset by software when the food tokens were acquired, and the battery was not actually replaced. As a result, it was confirmed that the body temperature was still appropriately controlled (Extended Data 4). Therefore, from these comparisons, it can be concluded that ENH’s motor control does indeed contribute to temperature control.

### Emergent Behaviours

To observe the overall behaviour acquired by ENH in more detail, we fixed some of ENH’s interoception to specific values and input them into the network, and observed ENH’s responses to each situation. First, we observed the qualitative behaviour of ENH at reduced values of the battery level; Fig. 4a shows an example of this behaviour, in which a series of navigation-like movements were observed to approach the food tokens. (Supplementary Video 2).

**Fig. 4.**
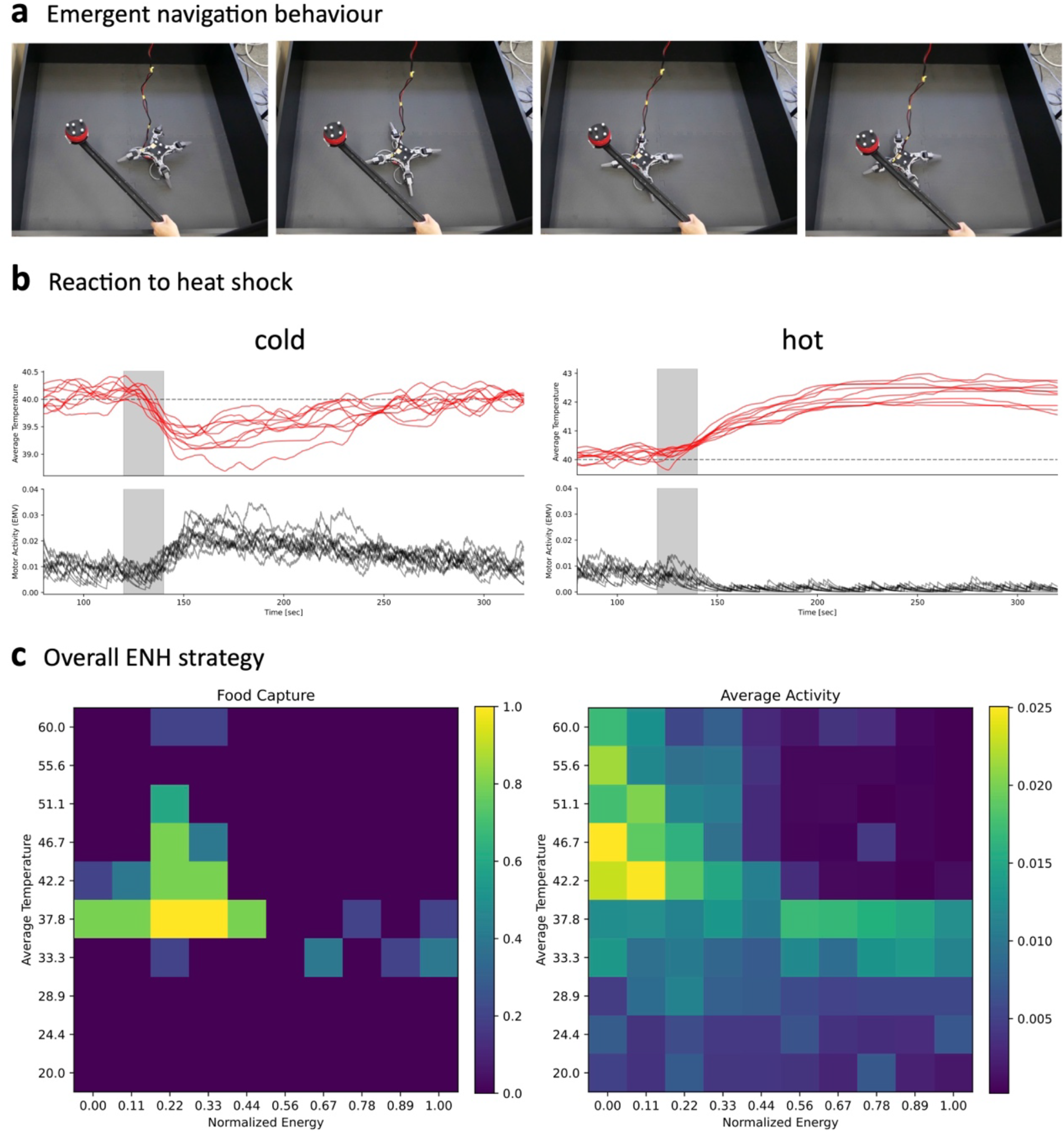
Emergent Behaviours. **a**, Navigation-like behaviour toward a food token observed when ENH’s observed battery charge is fixed at a low value. Although ENHs are not explicitly trained for navigation behaviour, a series of navigation-like movements are observed as a solution to homeostasis maintained by interactions with food tokens. **b**, Results of ENH response to thermal shocks. Results for both cooling (“cold”) and heating (“hot”) conditions are represented. The thermal shock is given to ENH only during the period of the grey background in the plot. The red line in the upper panel represents the body temperature (average motor temperature) and the dashed line represents the target temperature (40°C). The lower panel represents the time series of the corresponding intensity of motor activity. **c**, The overall behavioural strategy of ENH visualized by fixing our ENH’s interoception.

After setting the observed value of remaining battery charge large enough and confirming that ENH’s body temperature was stable enough, an external thermal shock was applied and the intensity of ENH’s movements was measured, defined as the sum of squares of the output signals to the motors. Fig. 4b shows the body temperature of the ENH (top) and the corresponding intensity of the movement (bottom) when a temperature shock was applied, measured over 10 trials. The left panel shows the response to the cooling shock and the right panel shows the results of the response to the heating shock. It is clear from the figure that in cooling, the intensity of movement of ENH temporarily increases after temporal decrease in body temperature, and the body temperature is restored to the target temperature (Supplementary Video 3). Conversely, in response to a heating shock, the intensity of ENH movements decreases to near zero, indicating that they are waiting for heat dissipation (Supplementary Video 4).

The overall picture of ENH behaviour depending on the internal state can be better understood in Fig. 4c. This panel shows the probability of food intake (left) and intensity of movement (right) when ENHs’ interoception were fixed at various values and their behaviour was recorded. As the heatmap shows, ENH took food tokens in the region of lower battery charge and increased exercise intensity in the region of lower body temperature. Furthermore, if we focus on the behaviour in the region where both body temperature and battery charge decrease in the heat map, we can confirm that ENH does not forage in this region and increases the intensity of movement. Therefore, it can be confirmed that the ENHs in this study have an overall strategy that prioritizes the control of body temperature before foraging.

## Conclusion

Homeostasis is a fundamental property of animals in nature, used to explain their behavior. However, how it produces integrated behaviours essential for survival has remained a mystery.

In this study, we utilized the physical dynamics of the robot’s internal state to optimize behavior, with homeostasis as the sole learning objective. Consequently, we demonstrated for the first time that integrated behaviours, including walking, foraging, and temperature control, could emerge in real-world robotic systems, termed the embodied neural homeostat.

The potential applications of our work are vast and diverse. For instance, developing highly adaptive systems that emulate the sophisticated integrated behaviours observed in nature. Using homeostasis as a principle, it might be possible to construct physiological models in real systems. Furthermore, our work opens new frontiers in pet robotics, where machines can synthesize emergent behaviours from their interactions with the physical environment and their embodiment, revolutionizing the concept of human-robot interaction.

## Supplementary Information

## Acknowledgement

I thank Jussi Saini of OTE Robotics for development support using RealAnt, Tatsuya Daikoku, Dongming Kim, Yoshia Abe, Washizu Shoya, Etsushi Arikawa, Yamato Shinomiya, and Emiri Ko for discussions. Fig. 1a was partly generated using Servier Medical Art, provided by Servier, licensed under a Creative Commons Attribution 3.0 unported license

## Author Contributions

N.Y. designed and implemented the reinforcement learning algorithm in *ENH*. N.Y. designed and implemented the robot system in *ENH*. N.Y. designed and implemented the evaluation framework for *ENH*. N.Y., H.K., Y.K. managed and advised on the project. N.Y. and H.K. wrote the paper.

## Methods

### Configuration of the robot system

The robot system used in the experiments was based on an off-the-shelf quadruped robot system, RealAnt [1] (OTE Robotics Ltd.), which has been reported to learn to walk using deep reinforcement learning in previous studies. It consists of eight Dynamixel AX-12A DC servo motors (ROBOTIS Ltd.), with two servo motors connected to each leg.

The servo motor was connected to the Arduino NANO 33 IoT via a connection board, and the Arduino was connected to an external PC via USB for serial communication, allowing the external PC to specify the target angle of servo motors and to obtain the current angle of servo motors, temperature inside the motors, and current target angle information.

The robot’s motor was powered by an external lithium-ion battery (Makita BL1415N). By connecting a power monitoring IC (INA226, Texas Instruments Inc.) to the battery’s power terminal, the battery’s current voltage and the amount of charge consumed [C] since the start of monitoring could be measured. The power monitoring IC was connected via I2C to the RaspberryPi 4 powered by the lithium-ion battery described above, and an external PC could observe the battery information via a wired connection to the RaspberryPi4. The lithium-ion battery outputs a voltage of around 14-16v, so a DC-DC converter was used to supply the appropriate voltage to each of the motor (12v) and RaspberryPi4 (5v).

In addition, markers attached to the robot and a food token (Fig. 1c, Food Token) were tracked by a motion capture system (Flex13, Acuity Inc.) consisting of six cameras to obtain their position and orientation information, which was used to constructs of the agent’s observations and recordings of the experiment. The robot’s controller was only given local information about the robot’s surroundings and not its global position or other information about itself. The agents could therefore leave the experimental environment, and to prevent this, the experimental environment was covered on all sides by wooden walls.

The observations input to the agent were composed of exteroception, proprioception and interoception respectively. For the agent’s exteroception, the position and posture information of the robot and the food token acquired by the motion capture system was used to construct a virtual 20-dimensional rangefinder output. The rangefinder returns the distance between the centre of the robot and the food token if the food token is within the 20-way angular region centred on the agent, otherwise it returns a value of zero. The maximum detection distance of the rangefinder was set to *d*_*max*_ = 1.5 [m]. The inputs to the agent were normalised by 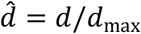 to have a value range of [0, 1] for each rangefinder output.

The agent’s proprioception consisted of a 32-dimensional vector connecting the time series of joint angles obtained at the last three times and the motor angle command values of the previous agent. This was intended to make the network robust to a slight time delay in the agent’s observation information.

We constructed the interoception as two-dimensional observation 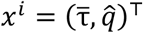: the first dimension uses the average value of the normalized temperature information provided by the servomotor 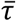(8-dimensional average motor temperatures) and the second dimension uses the normalised value of the charge consumption of the lithium-ion battery 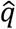. (normalised energy). Both of these changes depending on the dynamics of the physical quantities originating from the agent’s body, and the details of each value are given in the next section.

The agent’s output was an 8-dimensional vector representing target values of angles to the servomotor, with each axis normalised to [-1, +1]. The observation-action output cycle of the agent was set at one approximately 20 Hz.

### Average normalised motor temperature

The average motor temperature 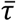 is the average of the temperature information (8 dimensions) provided by the servomotors of each joint. The temperature information τ_raw_ for each axis provided by the servomotors is in 1°C increments and is not smooth with respect to time. Therefore, the following smoothing was applied at each time step *t* to provide intermediate temperature information.

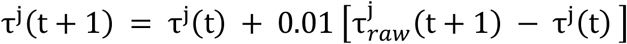

Where *j* is the index of the motor of the joint. The following linear transformation was used to map the range [20, 60] [°C] temperature to the range [-1, +1] when inputting to the network. This linear transformation maps 40°C to zero.

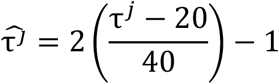

We used the average of these values for the interoception.

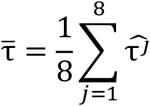

### Normalised Energy

The normalised energy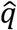. is the normalised amount of charge consumption *q*_raw_ [C] measured on the actual battery. We used the following transformation.

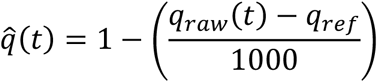

where *q*_*ref*_ is the charge consumption used as reference in the normalisation. Depending on whether the battery was replaced and how *q*_*ref*_ was updated, the following two resets were defined and used in the robot experiment.

#### Hard reset

The robot is switched off and the lithium-ion battery is replaced. The reference charge consumption is updated to the initial state (*q*_*ref*_ ≈ 0).

#### Soft reset

The robot remains switched on and stops the progress of the experiment. The reference charge consumption is updated to the charge consumption at the time *t*_*R*_ the soft reset is executed (*q*_*ref*_ = *q*_raw_(*t*_*R*_)).

### Configuration of simulators for training

To apply deep reinforcement learning for homeostasis with the Sim2Real approach, a multi-dynamics simulator is needed that includes not only the mechanical dynamics of the robot, but also the battery charge and motor temperature. In this study, each dynamics was modelled separately and this was constructed by fitting parameters based on data collected from a real robot.

Dynamics simulation of the robot body was created using the MuJoCo simulator [3]. The robot body and legs consist of a rectangular body and cylinders connected by rotary joints, respectively, and the mass and length of each part of the robot were measured by disassembling the actual robot and applying them to the model.

### Modelling thermal dynamics

A simple heat transfer model of the motor temperature was constructed for each motor independently. For the parameter optimisation described below, the following first-order Euler method was used for the simulation. For behavioural optimisation, a fourth-order Runge-Kutta method [4] was applied to the simulation using the same parameters. No significant differences in simulation results were found within the scope of this study.

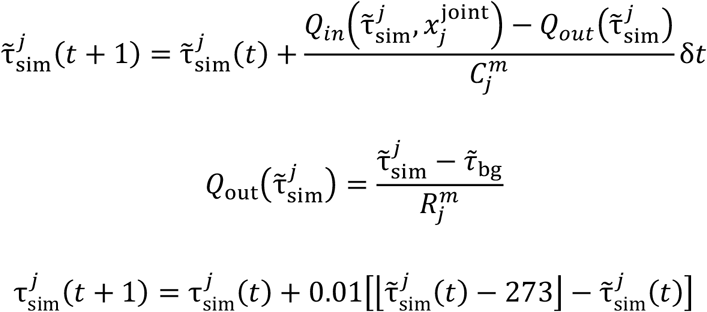

Where δ*t* is the time increment (=0.05 [s]), 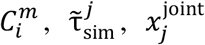 and 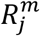 are the heat capacity, absolute temperature, joint angle, and thermal resistance. 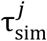 is the smoothed Celsius temperature time series and the floor function ⌊τ⌋ at update time returns the largest integer less than or equal to the real number τ. This reflects the fact that the temperature measurement of a real robot is given as an integer value. *Q*_out_ represents the heat flow due to the temperature difference between the motor and the outside, which is given by Newton’s law of cooling. 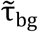 is the background temperature of the environment, eixed at 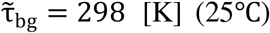. *Q*_in_ represents the heat flow rate from motor activity; The function giving *Q*_in_ was approximated by the heat flow model expressed below after trial and error.

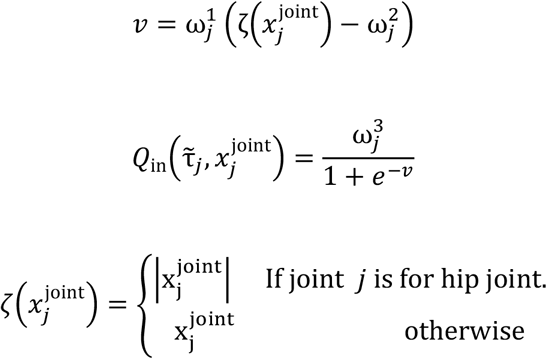

In equations, 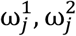, and 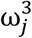 are the parameters of the heat flow model, respectively. For numerical stability, we replaced *v* by *v* = 100 sgn(*v*) when |*v*| > 100. Where sgn(⋅) is the sign function.

Thus, for each motor *j* the model has a 5-dimensional parameter of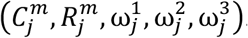.

To fit the parameters of the temperature model, the real robot was moved in this study and data was collected for 9,000 time periods (approximately 450 s) for each motor temperature along with joint angles. This data had intervals of 30 seconds each and repeated for two periods: first, a period during which the robot was moved with random behaviour selection at a constant intensity that varied with each interval; and second, a period during which the robot was kept in a default posture. In the first period, the following random action selections were applied to all joints.

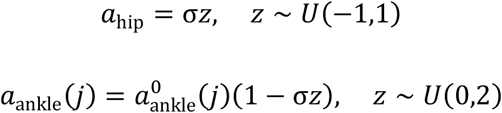

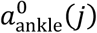 is the default posture value of the *j*th element of the leg-tip motor angle command value, specifically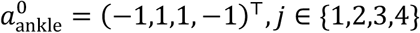. σ is the scaling parameter (movement intensity) for random behaviour that determines how far away from the default posture is allowed, with a value range of 0 < σ ≤ 1. In the actual sampling, the scaling parameter was sampled from a uniform distribution in the range [0, 1] at the start of the random behaviour period and the same value was used for that period. Two sets of data were prepared, one for training and one for testing, and used for later parameter fitting.

The fitting of the temperature model was performed independently for each motor and the parameter optimisation was performed by applying CMA-ES (Covariance matrix adaptation evolution strategy) [5]. The fitness function of CMA-ES was the function defined below.

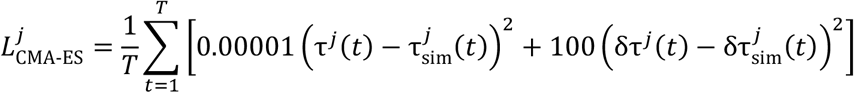

Where *T* = 9,000 and τ^*j*^ is the smoothed temperature of the motor *j* and 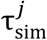 is the simulated temperature with the same joint angle input. δτ^*j*^ (*t*) and 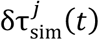 represent the temperature difference between the current time and the previous time. The implementation of CMA-ES is based on Optuna, which is available in the Python package [6]; the hyperparameters of CMA-ES were taken from Optuna’s default values, which were obtained after 1,000 trials of optimisation.

### Modelling of battery dynamics

In this study, a simplified battery dynamics model was adopted in which the normalised energy is consumed with constant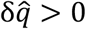.

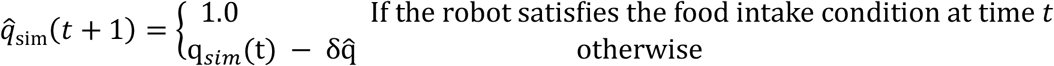

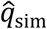 represents the normalised energy in the simulation. Food consumption in the simulation environment is determined when the centre position of the agent’s torso and the centre position of the food object are closer than 0.25 m.

To estimate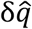, the amount of charge consumed was obtained by keeping the robot moving randomly at various movement intensities, similar to the random action selection described above. We confirmed that the above model fits well for a period of 5,000 steps of battery dynamics. Furthermore, by measuring the time variation of the charge consumption *q*_r*a*w_ for various vales of the movement intensity σ, the normalised energy reduction per step was estimated as 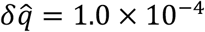 and used in subsequent simulations.

### Evaluation of real robot delays compared to simulations

Unlike a real robot, the robot in the simulator can obtain the joint angles of the robot without delay by directly referring to the values of the physics simulation. However, in a real system, there are differences between the actual joint angles and the physical situation due to errors in the model and the communication between the control PC and the robot board (Arduino). Therefore, in a preliminary experiment, the same joint angle commands were input to the simulator and the real robot system, and the amount of delay between the simulator and the real robot was evaluated by calculating the cosine similarity for a time series of joint angles of a certain length. As a result, the delay was approximately 0 to 1 step in the present experimental setup, and the delay was treated as negligible in the scope of the present experiment.

Reward setting based on homeostatic reinforcement learning

The only rewards used to train ENH were the following homeostatic reward settings using the agent’s interoception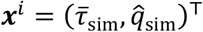.

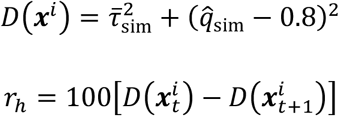

As indicated in the above equation, the target value of the normalised energy was set at 0.8.

### Network architectures

The network used in ENH consisted of a fully connected MLP with two hidden layers for both Actor and Critic (Extended Data 1). The hidden layers have a size of 256-64 dimensions and a GELU activation function was applied after layer normalisation; Actor uses a 9-dimensional, dimension-independent Beta distribution to represent the stochastic policy; due to the nature of the Beta distribution, each axis of Actor’s 9-dimensional output *a* outputs a value range of [0, 1] in each dimension.

The action *a* sampled from the policy is subsequently converted by Decoder into the actual final joint angle output of the robot: in Decoder, the 9th dimension output of *a* is considered as the parameter of the Bernoulli distribution, and the stochastic switching is performed by sampling the following.

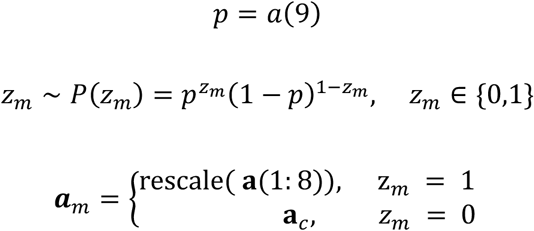

Where *a*(9) is the output of the 9th dimension of *a* and *a*(1: 8) is a vector consisting of the elements of the 1st to 8th dimensions of action *a* sampled from the policy. rescale(⋅) is a function that maps the output of the value range [0, 1] to the value range [-1, 1]. *a*_*c*_ = (0, −1,0,1,0,1,0, −1)^T^ is the posture in which all the robot’s motors are cooled, which we call the cooling posture. Referring to previous reinforcement learning studies with robots [7, 8], the behavioural output was smoothed by the following method to prevent damage to the robot’s body due to rapid changes in action.

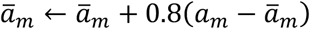

The ā_*m*_ calculated here is the final angle command to motors.

The reasons for the introduction of the Decoder, including the cooling posture, are explained below. From preliminary experiments, we found that the joint angles at which the motors do not generate heat and cooling of the robot’s body occurs are all within a very narrow range. For this reason, it was difficult to discover the cooling posture by probabilistic search in the present robot body, and it was necessary to explicitly incorporate the cooling posture. In a preliminary experiment, the agent was trained without the above decoder, and as a result, the agent was unable to find the cooling posture and an excessive increase in motor temperature was observed.

### Training conditions

Deep homeostatic reinforcement learning with PPO was used to train the agents. The optimisation was performed by 10 parallel sampling threads until the collection and training of samples of 3 × 10^8^ steps was completed. The hyperparameters of the PPO are described in Extended Data 2.

An environment reset was performed if the agent survived in the environment for 60,000 steps, or if the normalised energy was depleted 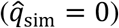, or if any of the eight dimensions of motor temperature exceeded the value range [20, 60]°C. Resetting the training environment involved randomly determining the initial position and orientation of the agent and the food position in the field. Also, after consumption of a food token, a new food token was generated at a random position in the field. In the test environment, the real robot was placed in the centre of the field with a defined orientation, reflecting the real robot experiment. The food tokens are generated in the same way as in the real robot process, where one of the four food positions is selected.

Domain randomisation was applied in the optimisation of the agent by randomly determining the physical parameters of the environment each time the training environment was reset. Domain randomisation was applied to the roughness of the floor surface (0-0.01 m), the coefficient of friction of the floor surface (1.3-1.7) and the mass and centre of gravity position of the agent by attaching random weights of 0-0.2 kg to its torso, which were updated to new values each time the environment was reset.

Training was performed with 20 different random number seeds. For application to the real robot, one of these 20 agents was selected and applied to the robot, which at the end of training had achieved 60,000 steps of survival in both the training and test environments.

### Experimental procedure on a real robot

There is one robot and one food token in the real environment. The internal states within the robot to be controlled are the average motor temperature and the normalised energy level. These vary according to the actual dynamics inside the body, which depend on the activity of the robot, respectively.

The entire experiment consists of a semi-automatic process where the procedure is directed by a PC, and the experimenter continues the experiment by replacing the batteries of the robot according to the instructions from the PC. The robot recovers its energy level by interacting with the food token in the environment. Specifically, the robot interacted with a food token and moved it 5 cm to indicate that the food had been “consumed”‘, and the experimenter performed a series of battery replacement sequences. The food token position was randomly selected from four different locations in the environment, and the next position was specified after the food token was obtained. However, the position within 30 cm of the robot’s centre position was not chosen, and the position was selected from the other three locations.

In the actual experiment, two lithium-ion batteries of the same type (Makita BL1415N) were used, and while one was in use, the other was charged and replaced in turn, thus allowing the experiment to proceed continuously. As charge consumption had to be sufficiently large to perform the charge due to the charger specifications, the experiment was continued with the same battery until the battery voltage fell below approximately 15.5 V (*soft reset*) and the battery was replaced when the condition for the first food consumption after the voltage fell below that level was fulfilled (*hard reset*).

### Effect of temperature drop due to hard reset

When the battery is replaced during the entire experimental process, the power supply to the robot itself is temporarily quiesced. This causes the temperature of the robot to drop during this time, which may assist the robot’s temperature homeostasis. Therefore, it was verified whether the robot’s temperature homeostasis could be maintained by soft reset only, without battery replacement.

Extended Data 4 shows the results of the verification from the fully charged state of the battery until the battery voltage dropped sufficiently to make the experiment unsustainable. By using soft reset only, the motor maintains the posture immediately after food consumption during the reset, which causes the motor to heat up. The figure shows that the temperature drop associated with a hard reset is not negligible, as the temperature drop is not observed, but we can confirm that the temperature of the agent can drop appropriately after food consumption and the body temperature is regulated even in the soft-reset only situation.

### Temperature variation in agents that do not include temperature homeostasis

The same training was used to realise the agent in the case of not including homeostasis with respect to the average motor temperature. The architecture of the agent in this case does not include a cooling posture.

This resulted in walking and navigation that achieved homeostasis of normalised energy even for the agent not containing temperature homeostasis, as shown in Extended Data 3. However, there is no control over the motor temperature, and it can be seen from the plots that the motor temperature continues to heat up. Therefore, if the robot does not control the motor temperature, the motor temperature will owner-heat and it is not possible to run the experiment for a long period of time.

### Observation of emergent behaviour

Behavioural observations by fixing interoception were applied to the ENH in the real robots in this study. By fixing the normalisation energy of the agent, the navigation and temperature-responsive behaviour of the agent was observed.

Supplementary Video 2 shows the agent’s navigation-like behaviour towards the food token observed when the normalisation energy was fixed to 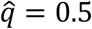. We can observe that the robot emerged the behaviour attracted to the object outside the body, even though the robot was optimised with the sole purpose of controlling the internal state of the body to a default value. This is because the dynamics inside the body are indirectly influenced by the action on the outside of the body, which can be interpreted as the complexity of the behaviour being generated not by the complexity of the reward function but by the complex dynamics created by the coupling of the dynamics inside and outside the body.

Supplementary Video 3 and Video 4 show the results of the change in the agent’s motor activity when the robot was exposed to a sudden temperature change with the normalised energy fixed at 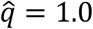, respectively. Motor activity was defined as the time-averaged exponentially weighted moving variance (EMV) using a smoothing factor of 0.01 for the RMS time series of ***a***_*m*_– ***a***_*c*_.

Both experiments consisted of the following three stages.

1. Place the robot in the environment for 120 s from the start of the experiment and wait until the robot’s motor temperature stabilises around 40 °C.
2. Apply a temperature shock for 20 s.
3. The robot’s response is then observed for 180 s.

The temperature shocks were evaluated for both situations: cooling of each motor with cold air and heating with warm air.

### Evaluation of overall behavioural characteristics dependent on interoception

For the evaluation in Fig. 3c, the behaviour was measured by fixing the ENH interoception to various values. In the measurements for this visualisation, the robot was initially placed in the centre of the environmental field and the food token was positioned 25 cm in front of the robot.

A single trial was terminated immediately after 10 seconds had elapsed or if the robot moved the food token 5 cm by physical interaction. The interoceptive was fixed at a specific value during the experiment, and in each situation the food token consumption event and the time-averaged intensity of the robot’s locomotion during the trial were measured. This measurement was performed on the inputs for each combination of values cut into a 10×10 mesh for the two-dimensional interoception space and five trials were performed for each combination and their means were obtained.

## Supplementary information

**Extended Data 1.**
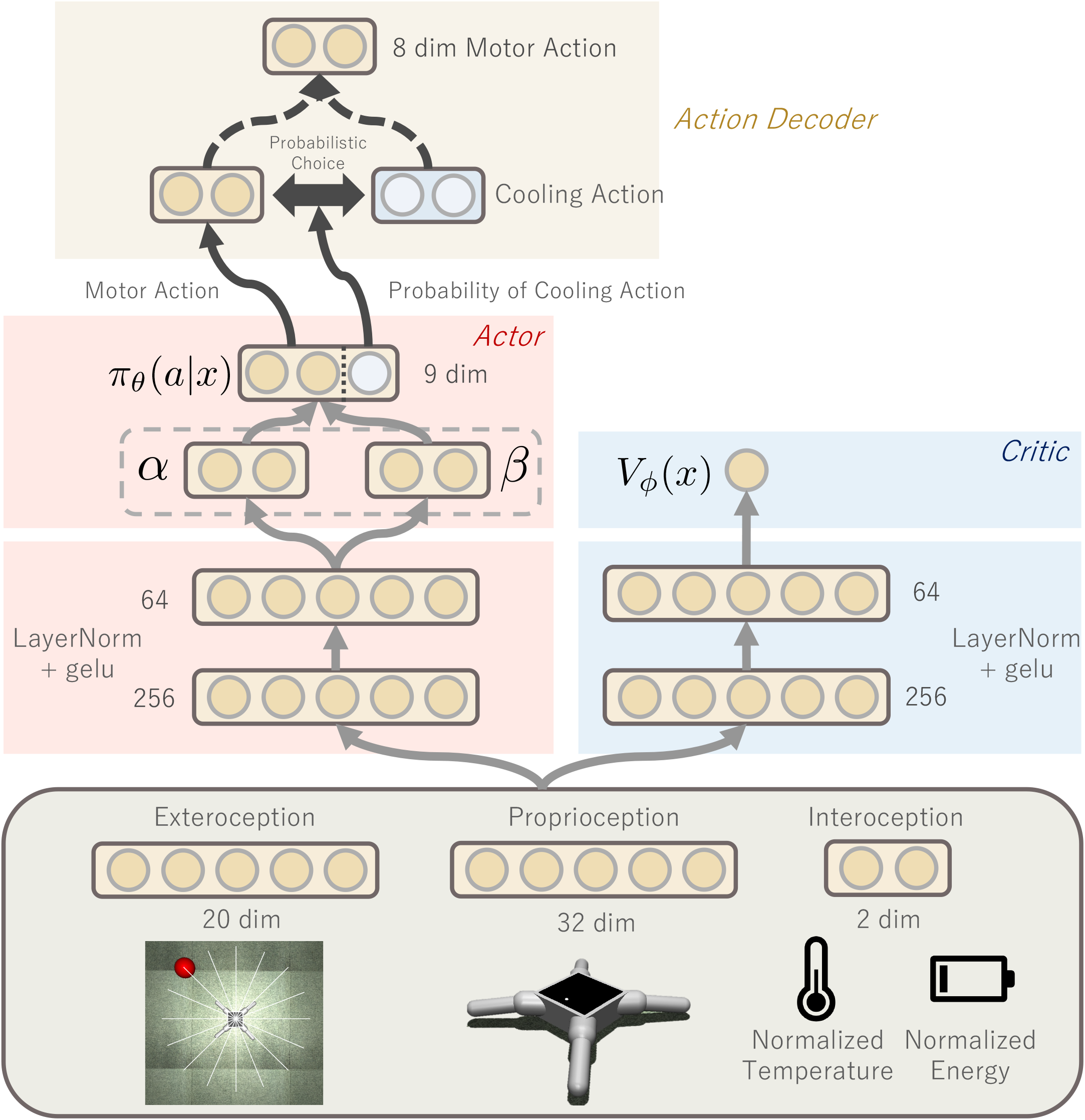

**Extended Data 2.**
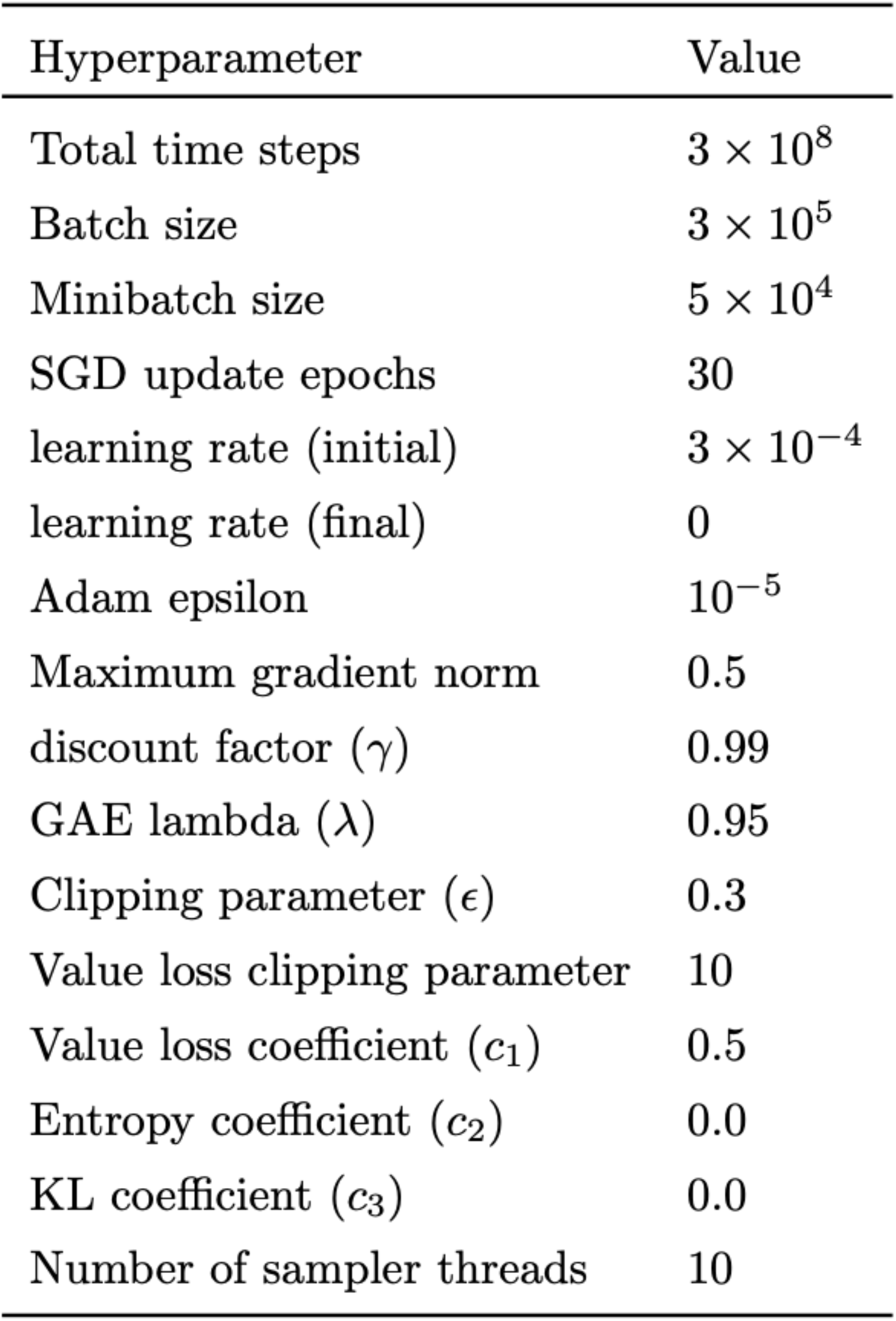
Hyperparameters of PPO

**Extended Data 3.**
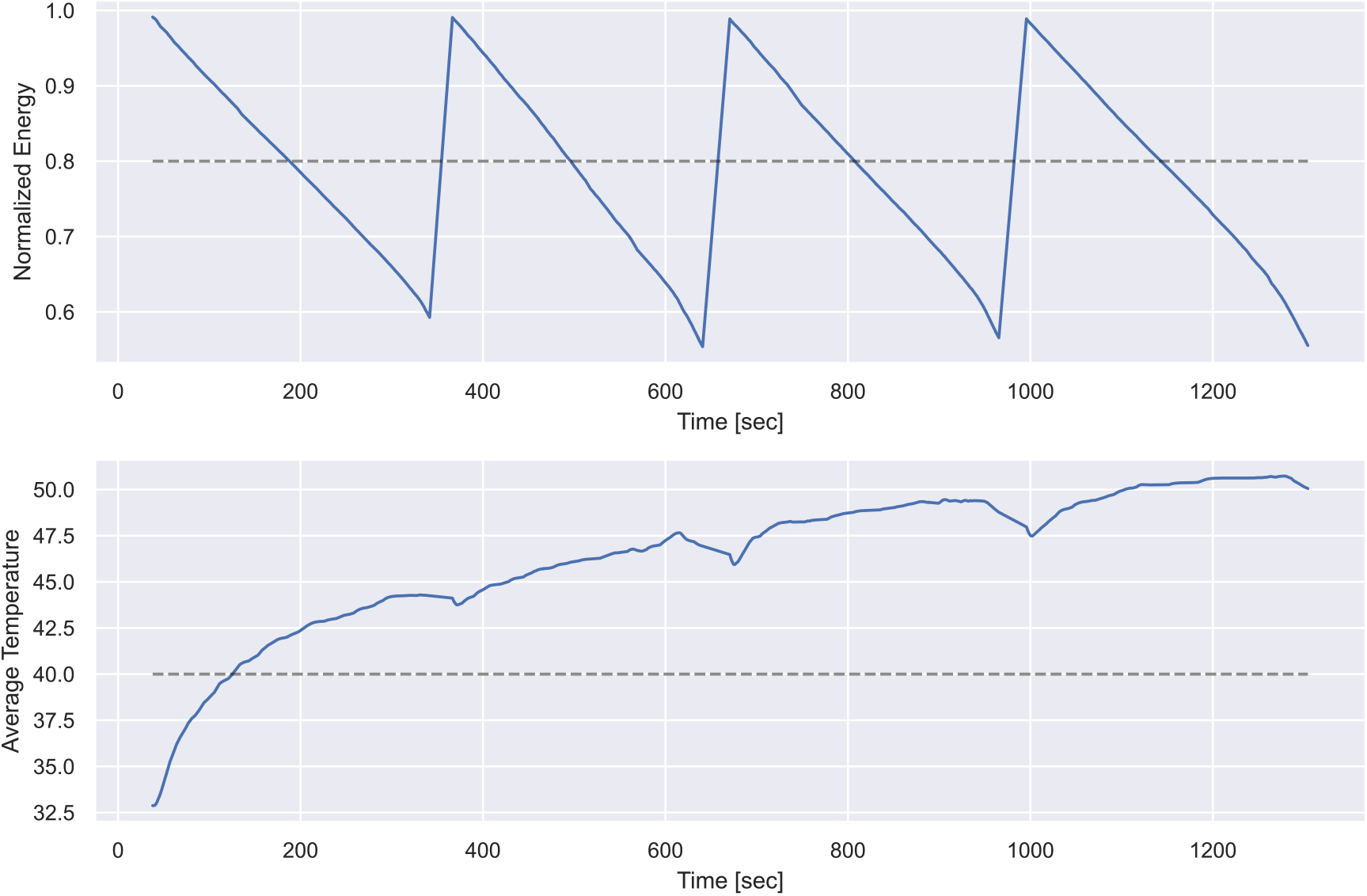
Results of a run under conditions where body temperature homeostasis is not a learning target.

**>Extended Data 4.**
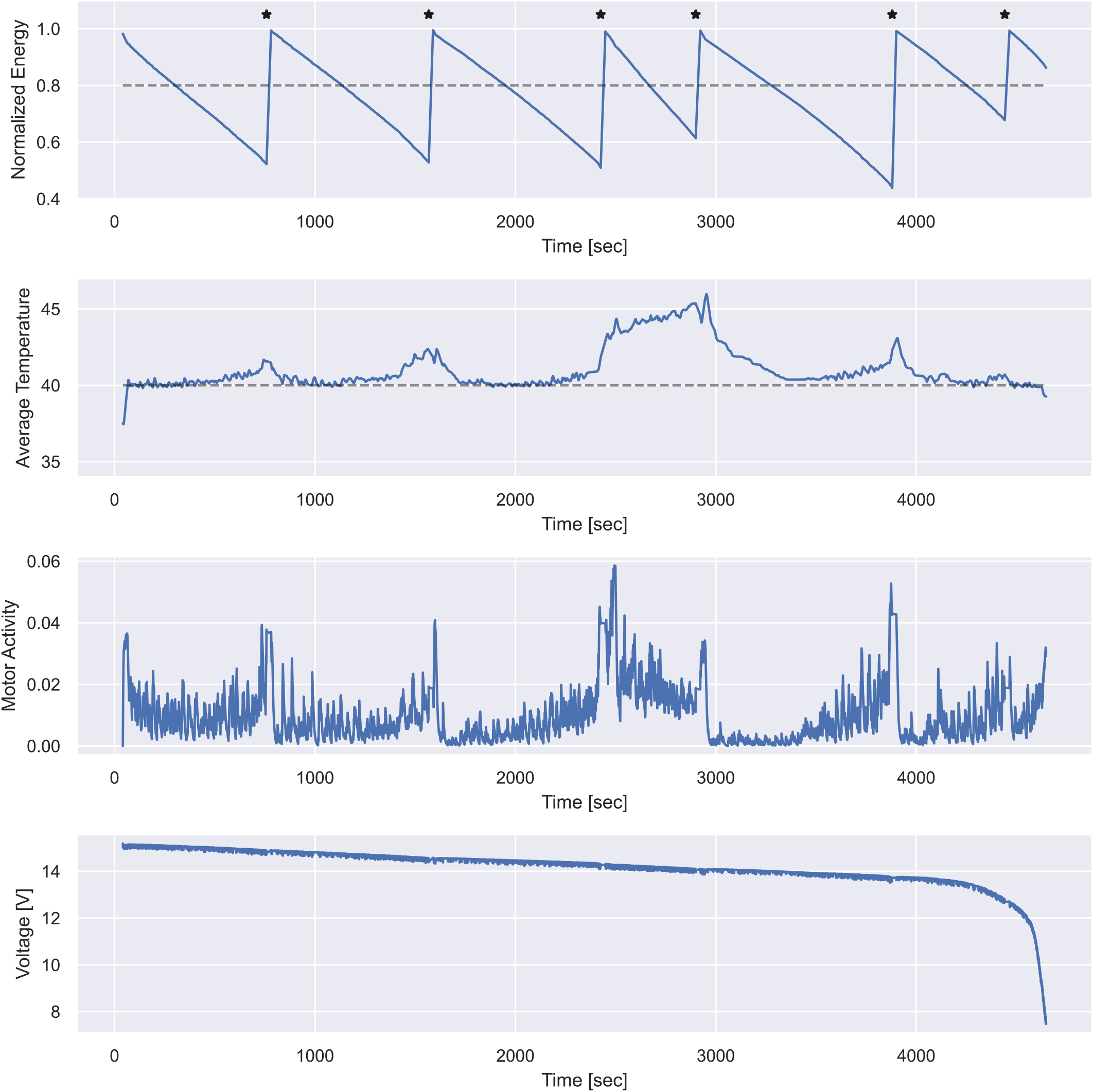
Results of a run with software simulated battery replacement.

**Video 1.**
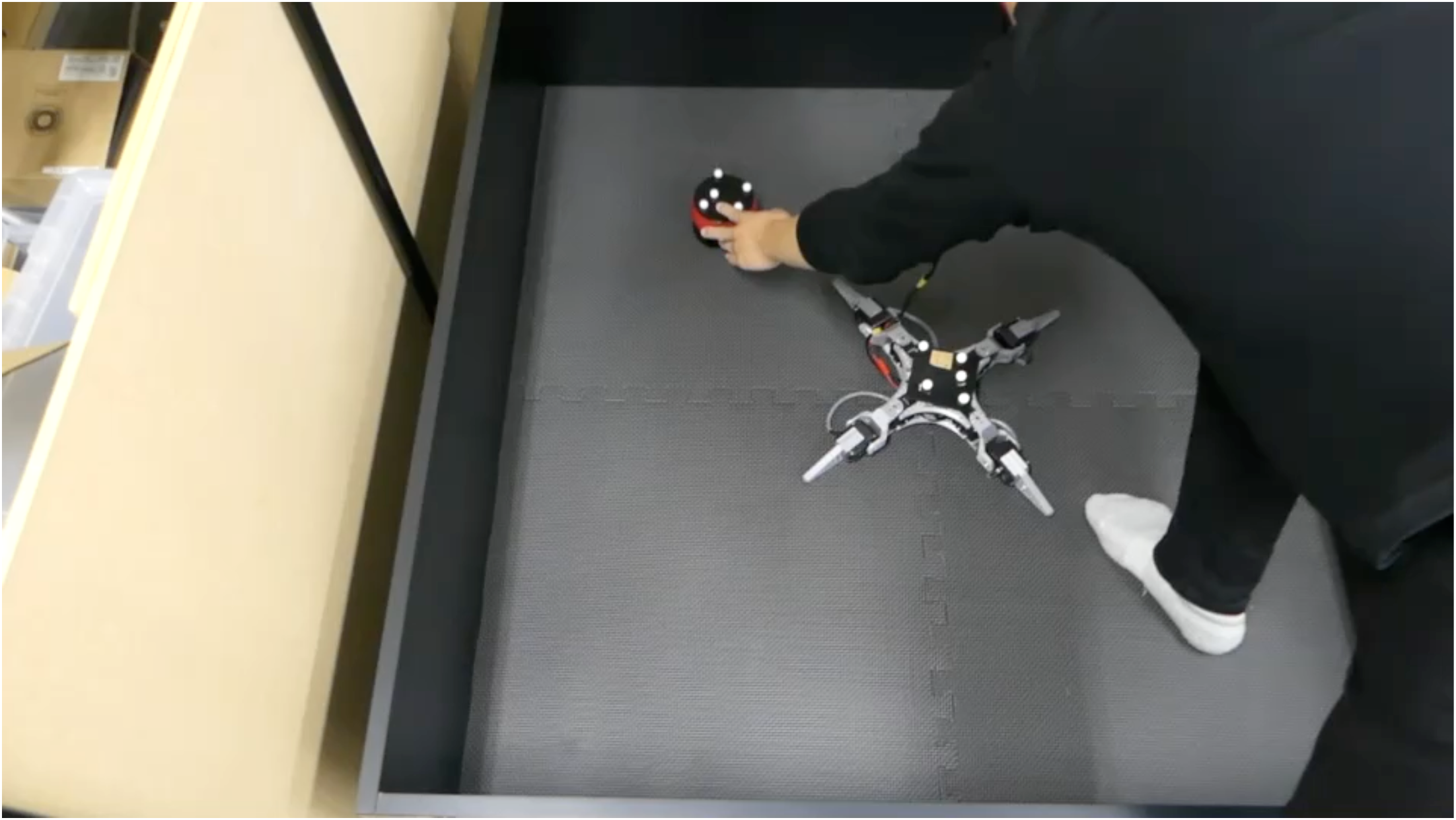
The demonstration of the whole experiment.

**Video 2.**
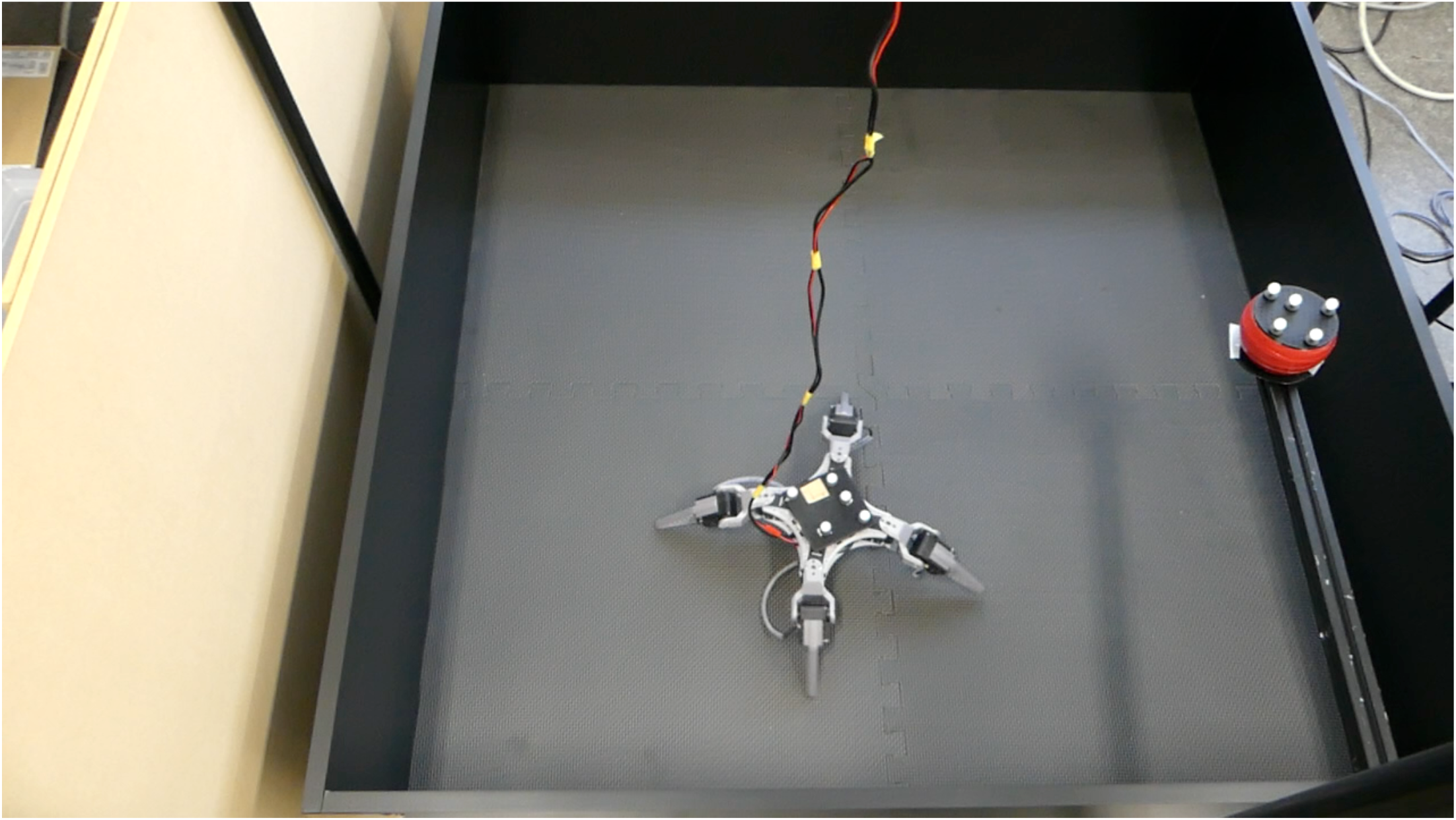
The demonstration of the emergence of navigation behaviour.

**Video 3.**
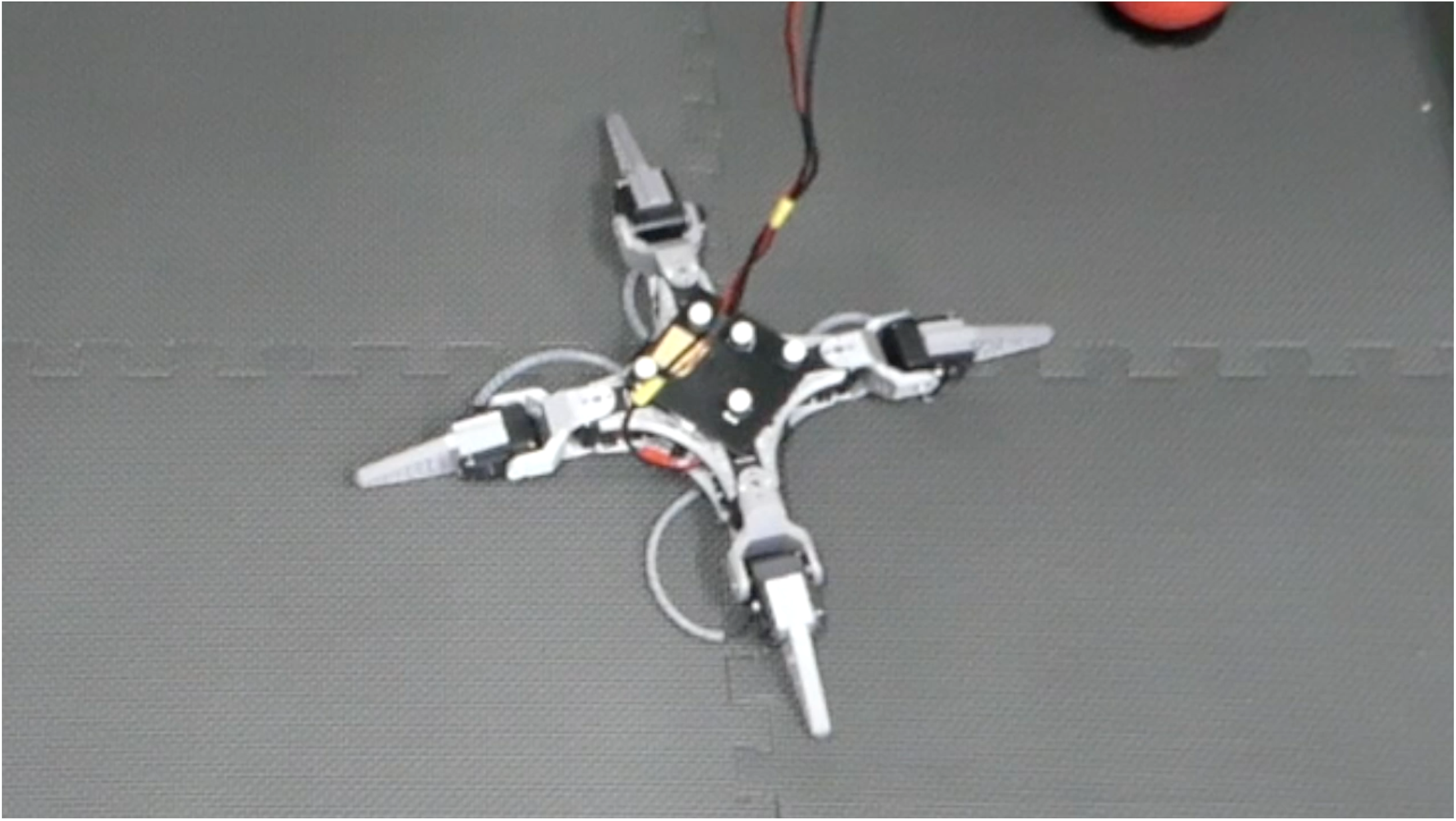
The demonstration of the cooling shock response.

**Video 4.**
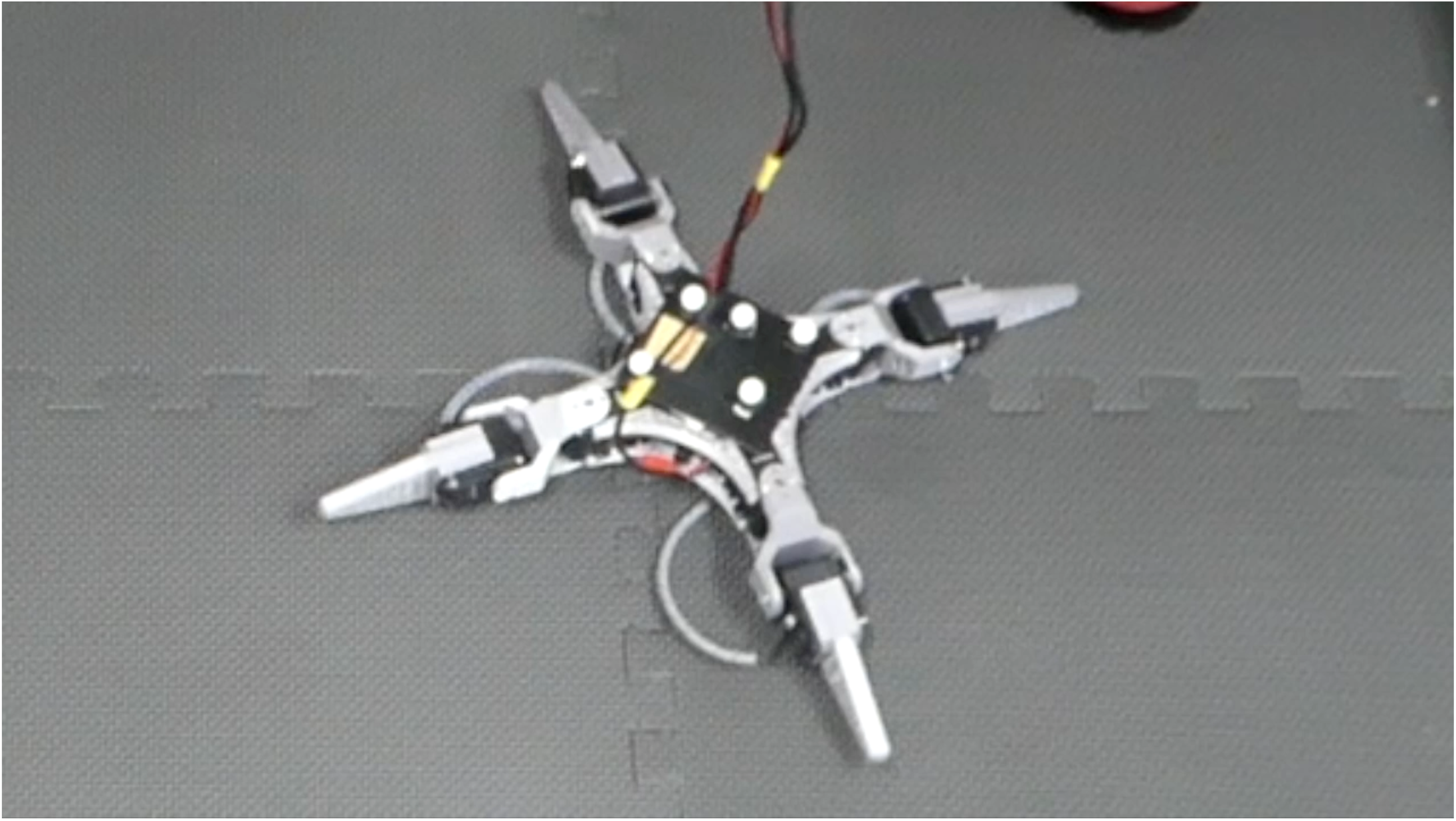
The demonstration of the heating shock response.

